# Ancient DNA Analysis of Archaeological Specimens Extends Chinook Salmon’s Known Historic Range to San Francisco Bay and Its Southernmost Watershed

**DOI:** 10.1101/2020.12.11.421065

**Authors:** Richard B. Lanman, Linda Hylkema, Cristie M. Boone, Brian Allée, Roger O. Castillo, Stephanie A. Moreno, Mary Faith Flores, Upuli DeSilva, Brittany Bingham, Brian M. Kemp

**Author notes:** Corresponding author. (R.B.L.).

## Abstract

Understanding a species’ historic range guides contemporary management and habitat restoration. Chinook salmon (*Oncorhynchus tshawytscha*) are an important commercial and recreational gamefish, but nine Chinook subspecies are federally threatened or endangered due to anthropomorphic impacts. Several San Francisco Bay Area streams and rivers currently host spawning Chinook populations, but government agencies consider these non-native hatchery strays. Using ichthyofaunal analysis of 17,288 fish specimens excavated from Native American middens at Mission Santa Clara circa 1781-1834 CE, 86 salmonid vertebrae were identified. Ancient DNA sequencing identified three of these as from Chinook salmon and the remainder from steelhead trout. These findings comprise the first physical evidence of the nativity of salmon to the Guadalupe River in San Jose, California, extending their historic range to include San Francisco Bay’s southernmost watershed.

**One Sentence Summary:** First application of ancient DNA to extend a species’ historic range finds Chinook salmon native to San Jose, California.

## Introduction

Chinook salmon (*Oncorhynchus tshawytscha*), also known as king salmon, are the largest salmon species in the world, and an important commercial and recreational gamefish. For the past several decades, Chinook have successfully spawned and reared in San Francisco Bay rivers and streams, a region where they are not considered native historically. Contemporary genetics studies suggest that the Bay Area’s spawning salmon are hatchery-derived fish, although a few individuals’ mitochondrial DNA fingerprints indicate that some are strays from wild stocks. Therefore, we applied ancient DNA (aDNA) sequencing of local archaeological fish samples to evaluate whether salmon historically utilized the Guadalupe River watershed in south San Francisco Bay, as establishing historic nativity could impact management decisions regarding the contemporary salmon population.

North American Chinook salmon currently range from Point Hope, Alaska (USA) to the Sacramento and San Joaquin Rivers in California’s Central Valley [1]. Nine different Chinook salmon evolutionary significant units (ESUs) are either federally threatened or endangered, with overall populations lingering at 1% or less than historic populations [2]. The California Central Valley fall-run Chinook salmon ESU, at the species’ southernmost limit, comprise 90% of recent California spawning fish, and traverse San Francisco Bay on their way to spawning in the Sacramento and San Joaquin Rivers’ inland watersheds [3]. However, government agencies do not recognize the San Francisco Bay’s coastal streams as historically utilized by Chinook salmon for spawning and rearing. Skinner’s extensive 1962 California Department of Fish and Game review opined that “although the fishery for king salmon is centered in the Bay Area, few kings actually spawn in any of the local streams”…and instead “pass through the Golden Gate to ascend the Sacramento and San Joaquin rivers on the way to ancestral spawning grounds in these rivers and their tributaries” [4]. Similarly, current National Oceanic and Atmospheric Administration (NOAA) National Marine Fisheries Sciences (NMFS) historic range maps show spawning and rearing habitat for Chinook salmon only in the Central Valley’s watersheds. The maps completely exclude San Francisco Bay’s coastal watersheds even though they are more proximal to the Pacific Ocean (see Fig 1) [5]. These reports on historical ranges are based on a paucity or absence of archaeological evidence, expert observer records, and museum records for Chinook salmon in either Bay Area streams, or in other streams further south on the Pacific Coast. However, for the last several decades, Chinook salmon have been documented spawning and rearing successfully in the lower, perennial reaches of larger Bay Area streams and rivers, including Walnut Creek and Napa River in the North Bay and the Guadalupe River in the South Bay [6–8].

**Fig 1.**
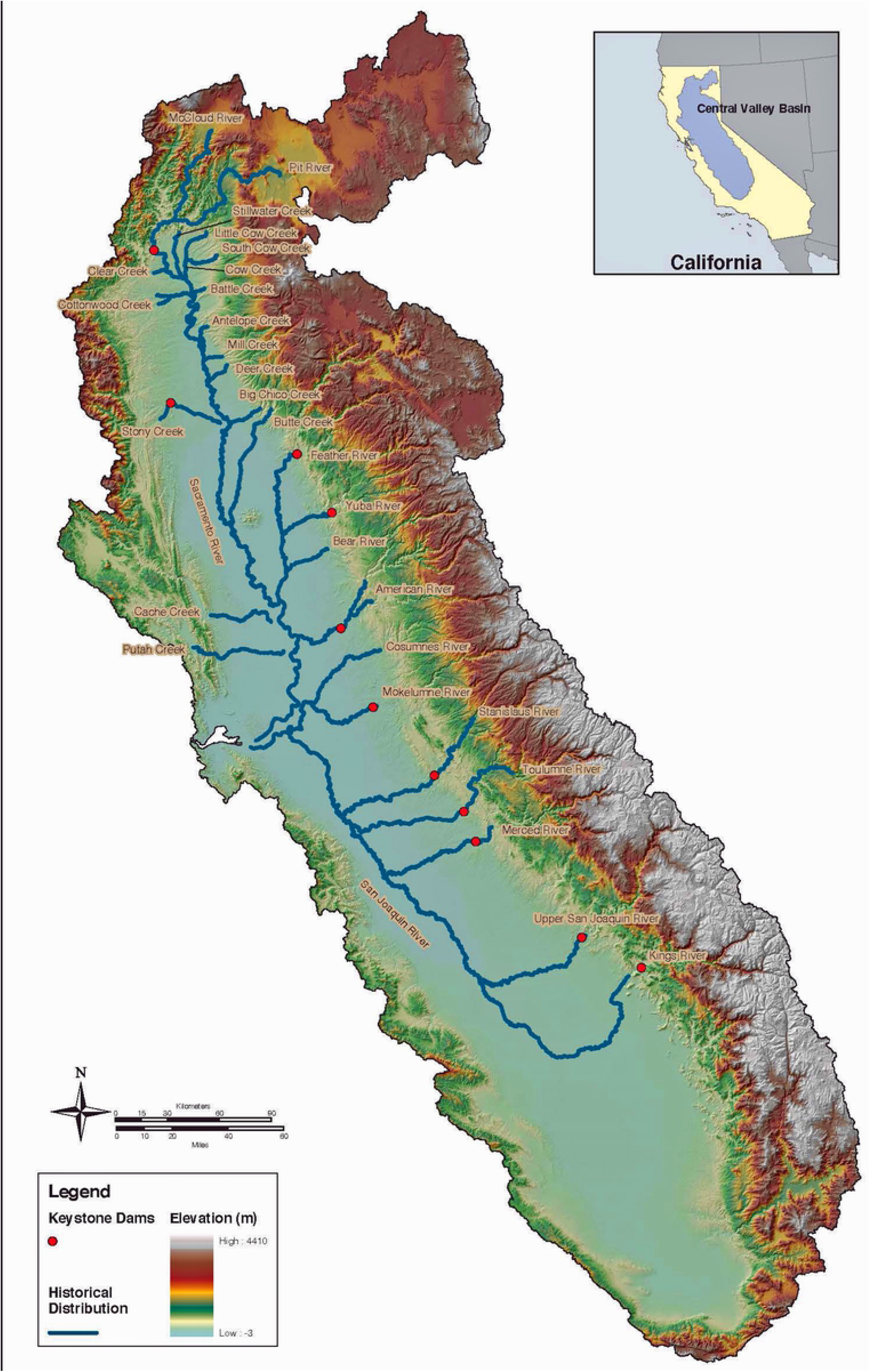
Historical distribution of Central Valley fall-run chinook salmon from NOAA Fisheries, Southwest Fisheries Science Center

The Guadalupe River hosts the southernmost of these nascent San Francisco Bay salmon runs. Salmon spawning has been observed in the river mainstem, which runs through San Jose, California (USA) at the extreme southern limit of San Francisco Bay, and its three tributaries: Los Gatos Creek, Guadalupe Creek, and Alamitos Creek. For several years, the South Bay Clean Creeks Coalition, a citizens-based watershed advocacy organization, has successfully monitored and conducted Geographic Information System (GIS) mapping of the carcasses of adult salmon and their redds (nests for egg deposition in stream gravels) in the Guadalupe River and its tributaries (see Fig 2). The salmon runs approached 1,000 adult fish in the late 1990s but were nearly extirpated in the early 2000s when the Army Corps of Engineers and Santa Clara Valley Water District constructed major anthropomorphic alterations to the river mainstem to mitigate flooding. San Francisco Bay watershed salmon runs have been attributed to hatchery strays, with very high rates of straying in recent drought years when hatchery-produced juvenile salmon were trucked downstream to the San Francisco Bay estuary to improve smolt survival [9]. Without a natal stream to home to as adults, these fish return and colonize new habitat for spawning.

**Fig 2.**
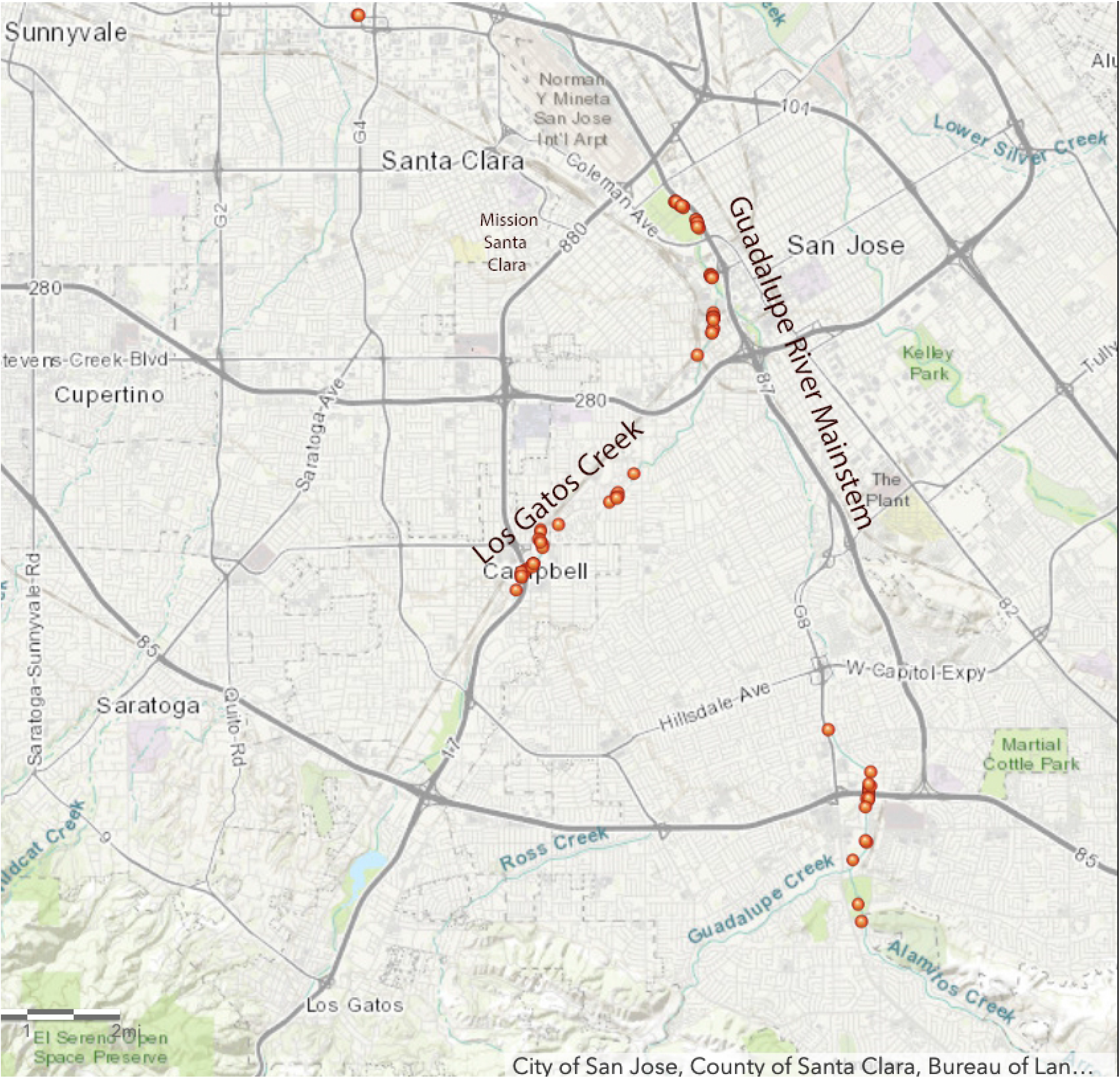
2019 Map of Chinook salmon adult carcass and redds in the Guadalupe River and its tributaries in San Jose, California. South Bay Clean Creeks Coalition January 2019

The only previously identified Chinook salmon archaeological specimens from the watersheds tributary to San Francisco Bay (defined as the estuary between the Sacramento and San Joaquin Rivers’ confluence and the Golden Gate strait) are present in Alameda Creek and Walnut Creek [10]. In addition, two historical observer records note Chinook salmon in San Francisco Bay’s coastal watersheds. First, an 1879 U.S. Commission on Fish and Fisheries report described obtaining Chinook salmon eggs in San Leandro Creek in the eastern Bay [11]. Second, a February 1904 newspaper account from San Jose, reported that “salmon and steelhead” were being “speared in local streams…even within the city limits” [12]. Two other expert accounts suggested that Chinook spawned in coastal watersheds even further south than San Francisco Bay, though neither collected actual specimens. In 1881, David Starr Jordan, ichthyologist and Stanford University’s first president, described Chinook spawning in coastal watersheds “From Ventura River northward to Behring’s [*sic*] Straits…” [13]; the Ventura River entering the Pacific Ocean 480 km (300 miles) south of San Jose. John Otterbein Snyder, another Stanford University ichthyologist, indicated in a 1912 report that Chinook salmon spawned in the Pajaro River, about 80 km (50 miles) south of San Jose [14].

Genetic studies of spawned out salmon carcass samples collected in the Guadalupe River watershed found that the majority of these fish are descended from inland Central Valley fall-run Chinook salmon hatchery stock [7,15,16]. However, two of these studies also identified some salmon with genetic profiles consistent with coastal rather than inland watersheds. The first, utilizing mitochondrial DNA, found two of nine haplotypes were unique to the Guadalupe River and two to the Russian River (California Coastal Chinook ESU) [15]. A more recent study utilizing microsatellite DNA found three of the 28 fish were more closely related to the Columbia River ESU [7]. Although these genetic studies suggested that today’s Guadalupe River salmon may not all be hatchery-derived, they do not resolve the question as to whether Chinook salmon historically utilized south San Francisco Bay streams.

The current study investigates this question by utilizing aDNA sequencing of salmonid vertebrae obtained via archaeological excavation at the historic site of the third Mission Santa Clara de Asís. This site, designated CA-SCL-30H, dates from 1781-1834 CE, and was located on the now-buried Mission Creek, a tributary of the Guadalupe River mainstem in San Jose. Although bone morphology can distinguish salmonids from other fishes, it cannot easily separate individual salmon species from one another, nor salmon (*Oncorhynchus* spp.) from rainbow/steelhead trout (*Oncorhynchus mykiss*) [17]. However, aDNA sequencing has been used to successfully differentiate each of the 12 species in the genus *Oncorhynchus* [18,19], potentially enabling us to ascertain whether any of the salmonid vertebrae from the excavation were Chinook salmon.

## Materials and Methods

### Museum Records Search for Chinook salmon in San Francisco Bay tributary streams

Ichthyology databases (FishNet2) and Integrated Digitized Biocollections (iDigBio) were searched for Chinook during the time period 1760-2020. No Chinook museum specimens were found for San Francisco Bay tributary streams, although multiple specimens were collected from the Bay itself. In addition, inquiries to individual museum curators found no Chinook salmon specimens for the above criteria at the California Academy of Sciences, Berkeley Museum of Vertebrate Zoology, Natural History Museum of Los Angeles County, San Diego Museum of Natural History, American Museum of Natural History, Harvard Museum of Comparative Zoology, or the Smithsonian National Museum of Natural History.

### Archaeological context

Archaeological excavation of the Native American Rancheria associated with Mission Santa Clara de Asís was conducted by Albion Environmental under the direction of Santa Clara University from 2012-2016. The study site is located 4.0 km (2.5 miles) from the Guadalupe River mainstem, on its historical Mission Creek tributary [20] in the city of Santa Clara just west of San Jose. The Mission’s Rancheria housed the locally indigenous Tamien-speaking Ohlone, as well as Bay Miwok and Delta Yokuts recruited from the broader San Francisco Bay Area [21]. The fish remains, which are the focus of this study, were recovered from two archaeological investigations (Franklin Block 448 and St. Clare) within the Rancheria, specifically from subterranean pits dug into the subsoil between adobe housing blocks [22]. The pits included numerous hearths, food processing tools, and food remains, indicating that these areas were used for the production and consumption of foodstuffs. Very fine-mesh 1.5 mm wet screens were used to isolate bony fish remains [23].

### Ancient DNA sequencing of salmonid vertebrae

Ancient DNA extracted from 86 vertebrae specimens identified as unspecified salmonids were sequenced at the Laboratories of Molecular Anthropology and Microbiome Research (LMAMR) ancient DNA laboratory at the University of Oklahoma to resolve specific *Oncorhynchus* species type using 148 bp sequences from the mitochondrial 12S gene (see Supplementary Materials for detailed methodology). Next, mitochondrial DNA from samples identified as Chinook salmon were sequenced along a different 563 bp stretch (from the end of the D-loop through tRNA-Phe and into the 12S gene) to gain additional phylogeographic information.

Further details on methods of ichthyofaunal analysis and ancient DNA sequencing are available in Supplementary Materials.

## Results

### Ichthyofaunal analysis

Fish remains were analyzed from multiple spatially distinct features at the Rancheria site, reaching a numeric total of 17,288 identifiable fish specimens (NISP). Freshwater fishes comprised 79-95% of the site assemblages by NISP, with most of the remaining specimens representing indeterminate freshwater/euryhaline species or euryhaline. Less than 1% of specimens were from marine fishes (Table 1). Of the 58 vertebrae identified as salmonid that also produced aDNA results, most of the vertebrae were very small, measuring ≤4 mm across the centrum diameter. Three specimens were all fragments but estimated at >10 mm or “large” (Table 2).

**Table 1.** Numbers of identified specimens (NISP) by habitat and two projects for fish remains at the Mission Santa Clara Rancheria archaeological site.

|  | Franklin Block Projects |  | St. Clare Project |  | Total |  |
| --- | --- | --- | --- | --- | --- | --- |
| Habitat | NISP | %NISP | NISP | %NISP | Total NISP | Total %NISP |
| Freshwater | 14,999 | 95.6% | 1,260 | 78.8% | **16,259** | **94.0%** |
| Indeterminate<br>freshwater/euryhaline | 202 | 1.3% | 32 | 2.0% | **234** | **1.4%** |
| Euryhaline | 437 | 2.8% | 305 | 19.1% | **742** | **4.3%** |
| Marine | 51 | 0.3% | 2 | 0.1% | **53** | **0.3%** |
| **Total** | **15,689** | **100.0%** | **1,599** | **100.0%** | **17,288** | **100.0%** |

**Table 2.** Species, archaeological context, and estimated vertebral centrum diameter for 58 salmonid vertebrae that produced aDNA results.

| Sample no. for<br>aDNA<sup>1</sup> | Archaeological<br>Feature<sup>2</sup> | Species ID by aDNA | Vertebra Size Estimate<sup>3</sup> |
| --- | --- | --- | --- |
| 2.4 | 155 | *O. mykiss* | 3 mm |
| 2.5 | 155 | *O. mykiss* | 3 mm |
| 2.6 | 155 | *O. mykiss* | 2-3 mm |
| 2.7 | 503 | *O. mykiss* | 1-2 mm |
| 3.1 | 79 | *O. mykiss* | 1-2 mm |
| 3.2 | 155 | *O. mykiss* | 2-3 mm |
| 3.4 | 503 | *O. tshawytscha* | >10 mm |
| 3.5 | 91 | *O. mykiss* | 4 mm |
| 3.6 | 155 | *O. tshawytscha* | >10 mm |
| 4.1 | 79 | *O. mykiss* | 2 mm |
| 4.3 | 119 | *O. mykiss* | 2 mm |
| 4.5 | 119 | *O. mykiss* | 3 mm |
| 4.6 | 119 | *O. mykiss* | 1-2 mm |
| 5.1 | 119 | *O. mykiss* | 2-3 mm |
| 5.3 | 119 | *O. mykiss* | 2 mm |
| 5.4 | 119 | *O. mykiss* | 2 mm |
| 5.5 | 119 | *O. mykiss* | 2-3 mm |
| 5.6 | 119 | *O. mykiss* | 1-2 mm |
| 5.7 | 119 | *O. mykiss* | 2-3 mm |
| 6.2 | 155 | *O. mykiss* | 2-3 mm |
| 6.3 | 155 | *O. mykiss* | 2-3 mm |
| 6.4 | 155 | *O. mykiss* | 2-3 mm |
| 6.5 | 155 | *O. mykiss* | 2-3 mm |
| 6.6 | 155 | *O. mykiss* | 2-3 mm |
| 6.7 | 503 | *O. mykiss* | 2 mm |
| 7.1 | 155 | *O. mykiss* | 2 mm |
| 7.2 | 155 | *O. mykiss* | 3 mm |
| 7.3 | 230 | *O. mykiss* | 1-2 mm |
| 7.4 | 503 | *O. tshawytscha* | Large |
| 8.1 | 66 | *O. mykiss* | 2-3 mm |
| 8.2 | 66 | *O. mykiss* | 2-3 mm |
| 8.3 | 66 | *O. mykiss* | 2-3 mm |
| 8.4 | 66 | *O. mykiss* | 2-3 mm |
| 8.5 | 79 | *O. mykiss* | 1-2 mm |
| 8.6 | 119 | *O. mykiss* | 2-3 mm |
| 9.1 | 119 | *O. mykiss* | 2-3 mm |
| 9.6 | 119 | *O. mykiss* | 2-3 mm |
| 10.1 | 119 | *O. mykiss* | 2-3 mm |
| 10.2 | 119 | *O. mykiss* | 2 mm |
| 10.3 | 119 | *O. mykiss* | 2 mm |
| 10.4 | 119 | *O. mykiss* | 2 mm |
| 10.5 | 119 | *O. mykiss* | 2 mm |
| 10.6 | 119 | *O. mykiss* | 2 mm |
| 10.7 | 119 | *O. mykiss* | 1-2 mm |
| 11.3 | 503 | *O. mykiss* | 2 mm |
| 11.5 | 155 | *O. mykiss* | 3 mm |
| 11.6 | 155 | *O. mykiss* | 3 mm |
| 11.7 | 155 | *O. mykiss* | 2-3 mm |
| 12.3 | 155 | *O. mykiss* | 2-3 mm |
| 12.4 | 155 | *O. mykiss* | 2-3 mm |
| 12.5 | 155 | *O. mykiss* | 3 mm |
| 12.6 | 155 | *O. mykiss* | 2-3 mm |
| 12.7 | 155 | *O. mykiss* | 2-3 mm |
| 13.1 | 155 | *O. mykiss* | 2-3 mm |
| 13.2 | 155 | *O. mykiss* | 2-3 mm |
| 13.3 | 155 | *O. mykiss* | 2-3 mm |
| 13.4 | 155 | *O. mykiss* | 3 mm |
| 13.5 | 155 | *O. mykiss* | 2-3 mm |

<sup>1</sup> 28 specimens did not produce aDNA results and are not included in this table. <sup>2</sup> Feature 503 is from the St. Clare project; all other features are from the Franklin Block projects. <sup>3</sup> Vertebral centrum diameter is an estimate based on pictures with scales. The two *O. tshawytscha* specimens with sizes >10 mm are based on approximately 1/4 of the centrum. The *O. tshawytscha* noted as "large" was heavily fragmented, but clearly from a large vertebra.

### Results of ancient DNA sequencing

Based on the 148 bp sequence of the mitochondrial 12S gene, all vertebrae with adequate DNA (58 of 86, or 67.4%) were identified as *Oncorhynchus* species, confirming the ichthyofaunal determination (Table 2). Of these 58 salmonid specimens, 55 were identified as rainbow/steelhead trout. Results for 53 of these 55 specimens were replicable. In the case of the other two samples, we were able to only produce results from a single PCR reaction. However, given their small size and the overall abundance of rainbow trout/steelhead found archaeologically, these identifications are likely correct. The remaining three specimens (sample numbers 3.4, 3.6, and 7.4) were identified as Chinook salmon (Table 2) and these results were replicable, and further validated by the additional mitochondrial DNA sequences described below. No fish DNA was observed in any negative controls. The 12S sequences were deposited at Genbank (Accession numbers MW086771-MW086828).

Specimens 3.4, 3.6, and 7.4 were additionally sequenced from nucleotide positions (nps) 570-1121 relative to a comparative full mitochondrial genome of Chinook salmon (Genbank accession number NC_002980.1). All of these sequences were replicable and have been deposited at Genbank (MW113717-MW113719). Further, they demonstrate the three samples as originating from three separate fish (see Supplementary Materials). These sequences make phylogenetic sense, in that they are closely related to other Chinook salmon sampled from the lower Pacific coast, and they further validate our species identification based on our mitochondrial 12S gene sequences.

## Discussion

We combined aDNA sequencing and ichthyofaunal analysis of archaeological samples to show that Chinook salmon were utilized by the indigenous peoples living at Mission Santa Clara de Asís in the 18^th^ and 19^th^ centuries. The results establish the first physical evidence that Chinook salmon spawned historically in a San Francisco Bay tributary watershed, the Guadalupe River. To the best of our knowledge, no other studies have combined these methodologies to extend the historic range of an animal species. One study utilized ancient DNA sequencing to confirm historical and ethnographic accounts of Chinook salmon and steelhead trout spawning as far upstream the Klamath River as the Upper Klamath Lake, but this finding did not extend the known historical range limits of these anadromous fishes [24]. The current study also enabled us to determine that the mitochondrial DNA from the three Chinook salmon vertebrae were from three different individual fish. This finding was consistent with one of the three fish coming from a different archaeological feature (Table 2).

Furthermore, the vertebrae identified as Chinook in our study comprised all three specimens from large individuals, indicating that these were adult fish that ascended the Guadalupe River watershed to spawn. Adult salmon are semelparous, dying after spawning, unlike steelhead which are iteroparous. The fish remains found in the Mission Rancheria could not be hatchery strays, as these samples significantly antedate the first salmon hatcheries established in California—in 1870 for trout and 1874 for salmon [25]. Although we cannot definitively exclude that the three Chinook salmon identified by aDNA were not trade items from other regions, multiple lines of evidence support the argument that these salmon specimens were locally caught. First, less than 1% of the of 17,000+ analyzed specimens were marine fishes, and many of the latter were species caught intertidally, indicating that local indigenous peoples were not exploiting pelagic marine fish species. Second, there are no historical accounts of Pacific coast salmon fishing by California tribes north of San Luis Obispo 298 km (185 mi) south of San Jose, although there is archaeological evidence of salmon take in northern San Francisco Bay and the Central Valley’s watersheds [26]. Third, adult salmon are large, averaging 13-14 kg (29-31 lbs) and may reach 59 kg (130 lbs), unwieldy for transport whole even over short distances. Instead, based on ethnographic descriptions of other Native Californian groups, large fish would more likely have been filleted to remove bones, then either dried into strips or pounded into fish flour for trade, and thus would have been devoid of large vertebrae [27,28].

Today, several San Francisco Bay coastal streams and rivers provide habitat that is suitable for Chinook salmon. Successful spawning and rearing have been documented in the Guadalupe River in the South Bay over the last four decades, and in a newspaper record from 1904 that appeared capable of differentiating steelhead trout from salmon [12]. Ethnographic researchers describe the Ohlone and most other Northern California indigenous people as catching steelhead trout and salmon efficiently by use of weirs or long nets stretched across streams (see Fig 3), and also via loop nets and harpoons or spears at rapids [27,29]. John Peabody Harrington, an ethnologist and linguistics expert who studied California’s indigenous peoples, specifically described the Ohlone people in the Santa Clara Valley as using spears and nets to fish (see Fig 3) [30]. Of interest, no fishhooks or spearheads were found in the Mission Santa Clara excavations, although fishing net weights were identified. Although the majority of Mission Rancheria residents were Ohlone, Bay Miwok, and Delta Yokuts peoples for whom there are no records of major salmon ceremonies [30], an analysis of West Coast tribes who utilized salmon extensively for food found no correlation with tribes who had salmon ceremonies versus those that did not [31].

**Fig 3.**
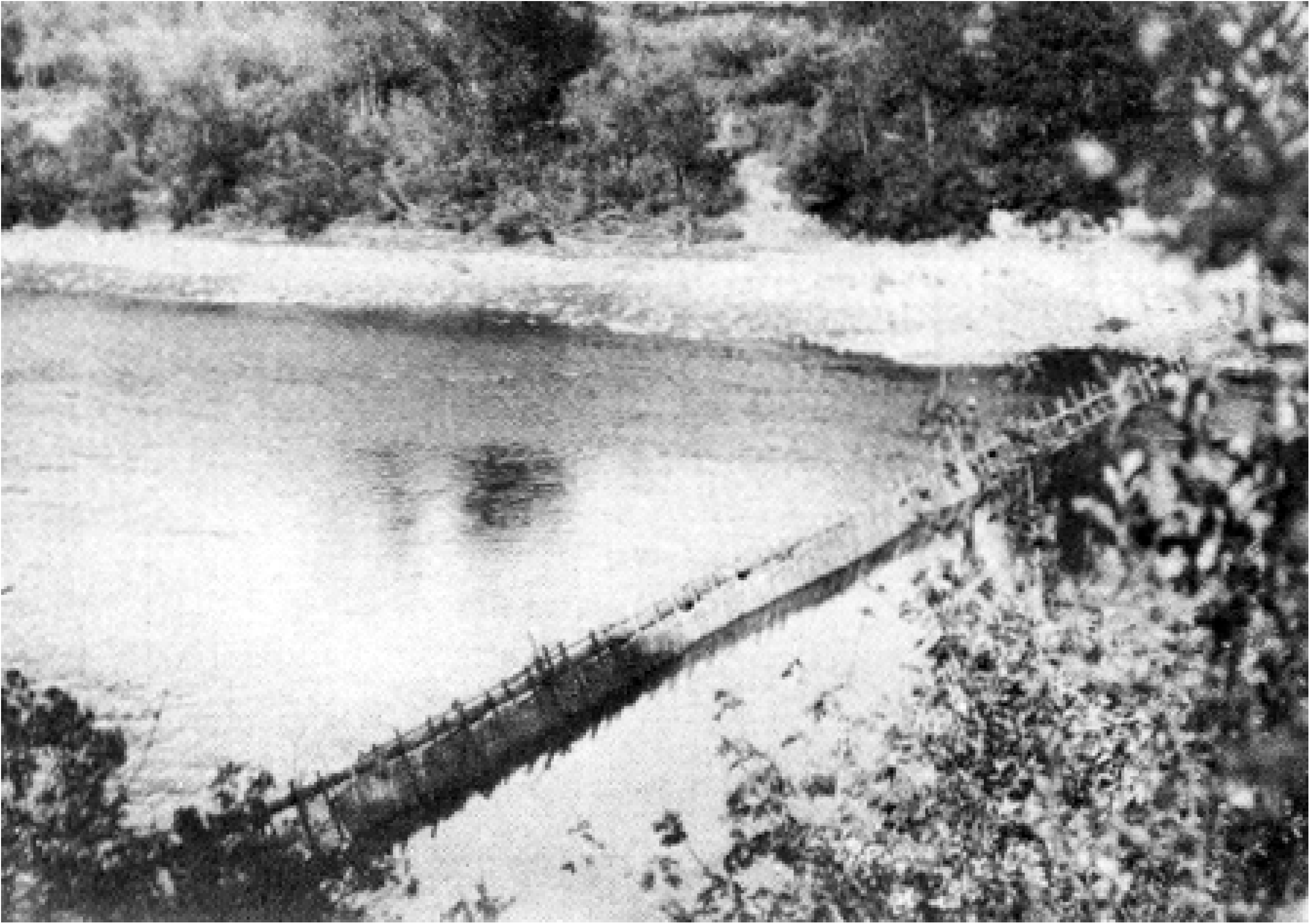
Hupa salmon weir on lower Trinity River, After P. E. Goddard, Life and Culture of the Hupa, University of California Publications American Archaeology and Ethnology 1, 1-88 (1903).

Why would there be apparent discordance between an absence of museum records and our finding of late18^th^/early 19^th^ century Chinook salmon in San Jose? First, only two California museums established ichthyology collections before 1900, the Stanford Museum and the California Academy of Sciences, and these were damaged or destroyed in the 1906 San Francisco earthquake and fire [32]. Second, our search of Chinook salmon museum records in the Bay Area revealed only fish caught in the bay itself. These specimens were likely most easily obtained from fish markets or from commercial fisherman targeting the large salmon runs traversing the bay en route to the Central Valley, rather than sourced from the smaller numbers of fish in the Bay’s tributary streams.

Daniel Pauly’s “shifting baselines syndrome” suggests that each generation of fisheries scientists accepts as a baseline the stock size and species composition that occurred at the beginning of their careers, and uses this to evaluate changes [33]. This phenomenon may be reflected in overly conservative governmental range maps for the southern limit of Chinook salmon. In fact, underestimation of the historic range for salmonids has occurred in at least three other North American examples. First, Chinook salmon were considered as nearly absent from the Russian River, part of the California Coastal Chinook ESU, until a 2007 publication of an underwater camera monitoring study documented an “abundant, widely distributed, and naturally self-sustaining Chinook population in the watershed” [34]. Second, a dearth of Atlantic salmon (*Salmo salar*) remains in the archaeological record were used to refute the nativity of Atlantic salmon to the coastal northeastern United States [35]. These assertions were overturned when Atlantic salmon fish scales were discovered in sediment cores taken from a nearshore pond in coastal New Jersey [36]. Third, based on the lack of archaeological findings in coastal streams south of San Francisco, lumber company affiliated biologists rejected Snyder’s determination of the southern limit of coho salmon (*Oncorhynchus kisutch*) at the San Lorenzo River in Santa Cruz County, California [14], and asserted that the coho salmon observed by Snyder in 1912 were hatchery strays [37]. However, one year later, archaeologists identified coho salmon vertebrae in Native American middens in nearby Año Nuevo State Park, reaffirming Snyder’s southern limit for coho [38]. These examples, coupled with the current study’s findings, should encourage California’s natural resource managers and scientists to question received wisdom as to the historic range of California’s fauna, and to apply new technologies, such as aDNA sequencing, to re-examine the evidence.

There are several limitations to the current study. Chinook salmon may have been present in the Guadalupe River watershed only intermittently, that is, only when conditions were favorable. All three of the Chinook salmon specimens date broadly to the Mission Period (1760-1834 CE) and while the specimen from the Franklin Block Feature 155 was from the Late Mission Period and could be differentiated spatially from the other two specimens from the St. Clare site, it could not be differentiated temporally from the others. Also, we cannot determine the size of the historic salmon population, which may have been low, or at minimum, highly variable over time in accordance with fluctuating climactic conditions. Lastly, our study does not verify whether today’s Chinook population is self-sustaining, that is whether local smolts eventually return as successfully spawning adults. This should be further investigated via isotopic analysis of contemporary adult salmon carcass otoliths, as they possess the same elemental composition as when the otoliths formed in early life, and are thus natal stream specific [9].

## Conclusion

The absence of evidence should not be equated with evidence of absence [39]. This study provides the first physical evidence that adult Chinook salmon historically spawned in the Guadalupe River watershed and extends the historic range of coastal Chinook further south than previously recognized. These results contrast with a paucity of archaeological, historical observer, and museum records. Ancient DNA sequencing of other archaeology specimens may refine our understanding of the historical range of other species. Whether today’s Guadalupe River salmon are hatchery strays or not is moot. As stated above, there is evidence that the recent Guadalupe River population has had at least some genetic introgression from Russian River and Columbia River stocks. If this watershed is managed to enable a self-sustaining coastal Chinook population at the very southern border of its range, these fish may represent an important genetic reservoir of fish buffered against changing climatic conditions, such as global warming [40], and may potentially counter collapsing diversity in this salmon species [41].

## Acknowledgments

The authors wish to acknowledge the encouragement of Kenneth Gobalet to study the possible Chinook salmon archaeology remains discovered at Mission Santa Clara. We also with to thank the South Bay Clean Creeks Coalition for its contemporary GIS mapping of salmon carcasses and redds in the Guadalupe River watershed.

## Author contributions

**Conceptualization:** Richard B. Lanman, Linda Hylkema, Cristie M. Boone, Brian M. Kemp

**Data curation:** Linda Hylkema, Cristie M. Boone, Brian M. Kemp

**Formal analysis:** Cristie M. Boone, Brian M. Kemp

**Funding acquisition:** Richard B. Lanman, Stephanie A. Moreno

**Investigation:** Linda Hylkema, Cristie M. Boone, Brian Alleé, Mary Faith Flores, Upuli DeSilva, Brittany Bingham, Brian M. Kemp

**Methodology:** Linda Hylkema, Cristie M. Boone, Brian M. Kemp

**Visualization:** Cristie M. Boonee, Brian M. Kemp, Richard B. Lanman

**Writing - original draft:** Richard B. Lanman

**Writing - reviewing and editing:** Richard B. Lanman, Linda Hylkema, Cristie M. Boone, Stephanie A. Moreno, Brian Alleé, Roger O. Castillo, Mary Faith Flores, Upuli DeSilva, Brittany Bingham, Brian M. Kemp

## Supporting information

See Supplementary Materials and Methods for further detailed supporting information, including Results of aDNA Amplification and Sequencing of the Three Chinook Salmon Samples, and Additional References (1-11).

## Data and materials availability

All sequence data generated in this study are deposited at GenBank as described herein. All data needed to evaluate the conclusions in the paper are present in the paper and/or the Supplementary Materials. Additional data related to this paper may be requested from the authors.

## References

1. Behnke R. Trout and Salmon of North America. New York, NY: Simon and Schuster; 2010.

2. Yoshiyama RM, Fisher FW, Moyle PB. Historical Abundance and Decline of Chinook Salmon in the Central Valley Region of California. North Am J Fish Manag. 1998;18: 487–521. doi:10.1577/1548-8675(1998)018<0487:HAADOC>2.0.CO;2

3. Munsch SH, Greene CM, Johnson RC, Satterthwaite WH, Imaki H, Brandes PL, et al. Science for integrative management of a diadromous fish stock: interdependencies of fisheries, flow, and habitat restoration. Can J Fish Aquat Sci. 2020; 1–18. doi:10.1139/cjfas-2020-0075

4. John E. Skinner. An Historical Review of the Fish and Wildlife Resources of the San Francisco Bay Area. Sacramento, California: The Resources Agency of California Department of Fish and Game Water Projects Branch; 1962 Jun p. 226. Report No.: Water Projects Branch Report No. 1.

5. Schick, Robert S., Edsall, Arwen L., Lindley, Steven T. Historical and Current Distribution of Pacific Salmonids in the Central Valley, CA. Santa Cruz, California: NOAA Fisheries, Southwest Fisheries Science Center; 2005 Feb p. 25. Report No.: NOAA-TM-NMFS-SWFSC-369. Available: https://swfsc.noaa.gov/publications/FED/00743.pdf

6. Leidy, Robert A. Ecology, Assemblage Structure, Distribution, and Status of Fishes in Streams Tributary to the San Francisco Estuary, California. San Francisco Estuary Institute; 2007 Apr p. 194. Report No.: SFEI Contribution No. 530. Available: https://www.sfei.org/sites/default/files/general_content/No530_Leidy_FullReport_revised_0.pdf

7. Garza, John Carlos, Pearse, Devon. Population genetics of Oncorhynchus mykiss in the Santa Clara Valley Region. Final Report to the Santa Clara Valley Water District (SCVWD). 2008 Mar. Available: https://pdfs.semanticscholar.org/68c6/e7a7b71d4c63b942a4f7db1c6e67c8b29b62.pdf

8. Garza, John Carlos, Crandall, Eric. Genetic Analysis of Chinook Salmon from the Napa River, California. NOAA Southwest Fisheries Science Center; 2013 Jul. Available: https://naparcd.org/wp-content/uploads/2014/10/NapaRiverChinookReport2013.pdf

9. Willmes M, Jacinto EE, Lewis LS, Fichman RA, Bess Z, Singer G, et al. Geochemical tools identify the origins of Chinook Salmon returning to a restored creek. Fisheries. 2020; fsh.10516. doi:10.1002/fsh.10516

10. Gobalet KW, Schulz PD, Wake TA, Siefkin N. Archaeological Perspectives on Native American Fisheries of California, with Emphasis on Steelhead and Salmon. Trans Am Fish Soc. 2004;133: 801–833. doi:10.1577/T02-084.1

11. Atkins, Charles G. On the distribution of schoodic salmon, in: Part V, Report of the Commissioner for 1877. Washington, D.C.: U.S. Commission of Fish and Fisheries, Government Printing Office; 1879 p. 832.

12. Anglers Rejoicing Over Recent Freshets. Good Trout Fishing is Now Insured in This County. Salmon and Steelheads Being Speared in Local Streams. San Jose Daily Mercury. 22 Feb 1904. Available: http://infoweb.newsbank.com/iw-search/we/HistArchive/?p_product=EANX&p_theme=ahnp&p_nbid=W5DH4ECMMTQ4NDYxNjY5MS4yNDg5MzY6MToxMjoxMjguMzYuNy4xMzg&p_docref=v2:1126156B6E3010F0@EANX-1140166AB440A9C8@2416533-1140166B5CCC3140@5

13. Jordan DS, Gilbert CH. Notes on the fishes of the Pacific coast of the United States. Proc U S Natl Mus. 1881;4: 29–70. doi:10.5479/si.00963801.4-191.29

14. Snyder, John Otterbein. The Fishes of the Streams Tributary to Monterey Bay, California. Bull Bur Fish. 1912;XXXII: 1–72.

15. Nielsen, Jennifer L. Salmon from the Sacramento-San Joaquin Basin and Guadalupe River 1992-1994. California Dept. of Fish and Game, Anadromous Fisheries Division, Sacramento; 1999. Report No.: CDFG Technical Report FG 2081 IF.

16. Garcia-Rossi, Dino, Hedgecock, Dennis. Provenance Analysis of Chinook Salmon *(Oncorhynchus tshawytscha)* in the Santa Clara Valley Watershed. Report to the Santa Clara Valley Water District. Bodega Marine Laboratory, University of California at Davis; 2002.

17. Moss ML, Judd KG, Kemp BM. Can salmonids (*Oncorhynchus* spp.) be identified to species using vertebral morphometrics? A test using ancient DNA from Coffman Cove, Alaska. J Archaeol Sci. 2014;41: 879–889. doi:10.1016/j.jas.2013.10.017

18. Jordan LG, Steele CA, Thorgaard GH. Universal mtDNA primers for species identification of degraded bony fish samples: Molecular Diagnostics and DNA Taxonomy. Mol Ecol Resour. 2010;10: 225–228. doi:10.1111/j.1755-0998.2009.02739.x

19. Grier C, Flanigan K, Winters M, Jordan LG, Lukowski S, Kemp BM. Using ancient DNA identification and osteometric measures of archaeological Pacific salmon vertebrae for reconstructing salmon fisheries and site seasonality at Dionisio Point, British Columbia. J Archaeol Sci. 2013;40: 544–555. doi:10.1016/j.jas.2012.07.013

20. Friend, N., Healey, C.T., Hunter, Thos., Smith, C.L. Historical Atlas Map Of Santa Clara County, California. Compiled, Drawn And Published From Personal Examinations And Surveys. Thompson & West, San Francisco, California; 1876. Available: https://searchworks.stanford.edu/view/10453189

21. Albion Environmental, Inc., David J. Powers and Associates, Inc. Master Cultural Resources Treatment Plan for the Santa Clara University 2020 Plan. Santa Clara, California: Santa Clara University; 2015 p. 241. Available: https://www.santaclaraca.gov/home/showdocument?id=19134

22. Peelo, S., Hylkema, L., Blount, C., Garlinghouse, T., Ellison, J., Boone, C. M., et al. The Indian Rancheria at Mission Santa Clara de Asis: Cultural Resources Mitigation for the Edward M. Dowd Art and Art History Building and Parking Structure, edited by S. Peelo and L. Hylkema. Santa Clara University Cultural Resource Management Heritage Stewardship Reports, L. Hylkema, general editor. Santa Clara, California: Santa Clara University; In Press.

23. Garlinghouse, T., Peelo, S., Blount, C., D’Oro, S., Brady, R., Ellison, J., et al. Archaeological Data Recovery for the St. Clare Residence Hall Storm Drain Project. Report Prepared for Santa Clara University. Albion Environmental, Inc., Santa Cruz, California; 2018.

24. Stevenson, Alexander E. Using Archaeological Fish Remains to Determine the Native Status of Anadromous Salmonids in the Upper Klamath Basin (Oregon, USA) Through mtDNA and Geochemical Analysis. Master’s Thesis, Portland State University. 2011. Available: https://doi.org/10.15760/etd.444

25. Leitritz, Earl. A History of California’s Fish Hatcheries 1870-1960. State of California The Resources Agency Department of Fish and Game; 1970 pp. 1–86. Report No.: Fish Bulletin 150. Available: http://content.cdlib.org/view?docId=kt5k4004bd&brand=calisphere&doc.view=entire_text

26. Baumhoff, Martin A. Ecological Determinants of Aboriginal California Populations. Univ Calif Publ Am Archaeol Ethnol. 1963;49: 155–236.

27. Rostlund, Erhard. Freshwater Fish and Fishing of Native North America. Berkeley, California: University of California Press; 1952.

28. Yoshiyama RM. A History of Salmon and People in the Central Valley Region of California. Rev Fish Sci. 1999;7: 197–239. doi:10.1080/10641269908951361

29. Swezey, S. L., Heizer, R. F. Ritual Management of Salmonid Fish Resources in California. J Calif Anthropol. 1977;4: 1–29.

30. Harrington, John P. Culture Element Distributions: XIX, Central California Coast. Univ Calif Archaeol Rec. 1942;7: 1–46.

31. Gunther E. An Analysis of the First Salmon Ceremony. Am Anthropol. 1926;28: 605–617.

32. Lanman, Richard B., Perryman, Heidi, Dolman, Brock, James, Charles D. The historical range of beaver in the Sierra Nevada: a review of the evidence. Calif Fish Game. 2012;98: 65–80.

33. Pauly D. Anecdotes and the shifting baseline syndrome of fisheries. Trends Ecol Evol. 1995;10: 430. doi:10.1016/S0169-5347(00)89171-5

34. Chase, Shawn D., Manning, David J., Cook, David G., White, Sean K. Historic Accounts, Recent Abundance, and Current Distribution of Threatened Chinook Salmon in the Russian River, California. Calif Fish Game. 2007;93: 130–148.

35. Carlson, Catherine C. Where’s the salmon? A reevaluation of the role of anadromous fisheries in aboriginal New England in Holocene human ecology. In: Holocene human ecology in northeastern North America. George P. Nicholas, Northeastern Anthropological Association, editors. New York: Plenum Press; 1988.

36. Daniels RA, Peteet D. Fish Scale Evidence for Rapid Post-Glacial Colonization of an Atlantic Coastal Pond. Glob Ecol Biogeogr Lett. 1998;7: 467–476. doi:10.2307/2997716

37. Kaczynski VW, Alvarado F. Assessment of the Southern Range Limit of North American Coho Salmon. Fisheries. 2006;31: 374–391. doi:10.1577/1548-8446(2006)31[374:AOTSRL]2.0.CO;2

38. Adams PB, Botsford LW, Gobalet KW, Leidy RA, McEwan DR, Moyle PB, et al. Coho Salmon are Native South of San Francisco Bay: A Reexamination of North American Coho Salmon’s Southern Range Limit. Fisheries. 2007;32: 441–451. doi:10.1577/1548-8446(2007)32[441:CSANSO]2.0.CO;2

39. Rees MJ. From here to infinity: scientific horizons. London: Profile Books; 2011.

40. Keefer ML, Caudill CC. Homing and straying by anadromous salmonids: a review of mechanisms and rates. Rev Fish Biol Fish. 2014;24: 333–368. doi:10.1007/s11160-013-9334-6

41. Carlson SM, Satterthwaite WH. Weakened portfolio effect in a collapsed salmon population complex. Fleming IA, editor. Can J Fish Aquat Sci. 2011;68: 1579–1589. doi:10.1139/f2011-084

